# Common Ground in Chaos: Diversified Photodynamic Treatments Converge on a Unified Stress Architecture in *Escherichia coli*

**DOI:** 10.64898/2026.08.13.744726

**Authors:** Natalia Burzyńska-Młotkowska, Anna Wróblewska, Michał Wojciech Szcześniak, Mariusz Grinholc

## Abstract

The rise of antimicrobial resistance has intensified interest in antimicrobial photodynamic inactivation (aPDI) and antimicrobial blue light (aBL) as alternatives or adjuvants to conventional antibiotics. However, whether chemically distinct photodynamic treatments elicit a shared bacterial response remains unclear. Here, we integrated transcriptomic profiles of *Escherichia coli* BW25113 exposed to five short-term, sub-lethal photodynamic treatments: antimicrobial blue light (aBL), aBL combined with 5-aminolevulinic acid (aBL+ALA), rose bengal (RB), new methylene blue (NMB), and the cationic porphyrin TMPyP. Intersection analysis identified 891 conserved core genes differentially expressed across all treatments, of which approximately 98% changed in a consistent direction despite differences in photosensitizer chemistry and activating wavelength. Random-effects meta-analysis and robust rank aggregation prioritized 88 high-confidence genes, revealing induction of envelope stress and cytoplasmic protein quality control pathways alongside repression of acid resistance, hydrogen metabolism, molybdate transport, and biofilm formation. Regulon enrichment indicated that heat-shock sigma factor σ^32^/RpoH and the envelope-stress regulators CpxR, BaeR, σ^24^/RpoE, and PspF were enriched among induced genes, whereas GadW/GadX/GadE, Fur, and σ^38^/RpoS were enriched among repressed genes. Functional validation using selected single-gene Keio knockouts confirmed that deletion of conserved-core genes sensitized *E. coli* to photodynamic treatment and delayed post-treatment recovery in a modality-dependent manner. Moreover, RT-qPCR analysis of selected transcriptional responses confirmed the direction and overall pattern of RNA-seq-derived expression changes. Together, these findings define a unified conserved early survival program in *E. coli* after chemically distinct photodynamic treatments and identify stress-response modules that may serve as targets for potentiating aPDI.

**Graphical abstract:** 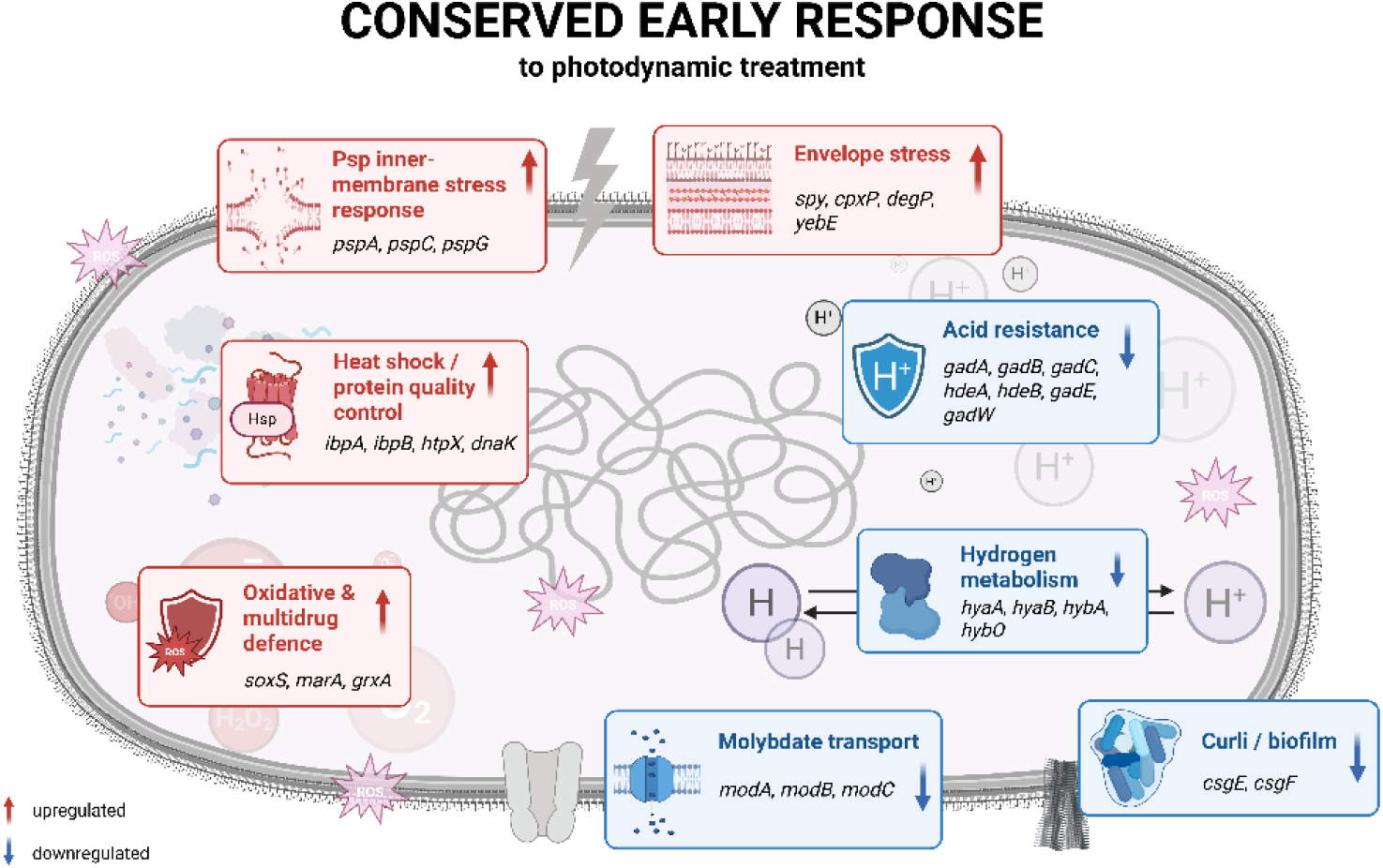

## 1. Introduction

The global rise of antimicrobial resistance (AMR) is among the most pressing threats to human health. In 2021, bacterial AMR was associated with an estimated 4.71 million deaths worldwide, of which 1.14 million were directly attributable to drug-resistant infections, with *Escherichia coli* remaining among the leading bacterial pathogens contributing to the global AMR burden [1]. The continuing rise in resistance, together with the limited innovation and pathogen coverage of the current antibacterial development pipeline, underscores the need for antimicrobial strategies that act through mechanisms distinct from those of conventional antibiotics [1], [2].

Antimicrobial photodynamic inactivation (aPDI) and antimicrobial blue light (aBL) are multi-target antimicrobial approaches considered to have a lower potential for resistance development than conventional antibiotics. Both rely on the light-driven excitation of photoactive molecules in the presence of molecular oxygen, resulting in the generation of reactive oxygen species (ROS) that simultaneously damage multiple bacterial targets [3], [4]. In aPDI, exogenous photosensitizers are activated at wavelengths matching their absorption spectra [5], whereas aBL excites endogenous bacterial chromophores, particularly porphyrins absorbing within the 400–470 nm range, and therefore does not require the addition of an exogenous photosensitizer [6]. Upon illumination, the excited photosensitizer can undergo Type I reactions, in which electron transfer results in the formation of superoxide (0_2_•^-^), hydrogen peroxide (H_2_0_2_), and hydroxyl radicals (HO•), or Type II reactions, in which energy transfer to molecular oxygen produces highly reactive singlet oxygen (¹O₂) [7]. Because photodynamic inactivation causes rapid oxidative damage to multiple cellular targets rather than inhibition of a single molecular pathway, the development of stable resistance is considered less likely than for conventional antibiotics, including after repeated cycles of sub-lethal exposure [8], [9], [10], [11].

However, the clinical translation of aPDI and aBL is limited by the heterogeneity of photosensitizers, light sources and treatment regimens, as well as by an incomplete understanding of bacterial responses to photodynamic stress. Transcriptomic studies have generally focused on a single photosensitizer, treatment modality, or time point, with limited comparison across chemically distinct approaches. It therefore remains unclear whether photodynamic treatments differing in photosensitizer chemistry and activating wavelength converge on a shared bacterial transcriptional response or instead elicit predominantly treatment-specific programs. Resolving this question may reveal conserved components of the bacterial stress response that could serve as modality-independent targets for potentiating photodynamic antimicrobial treatments.

In a previous study, we used time-resolved global transcriptomics to map how *E. coli* BW25113 reprograms gene expression in response to five photodynamic treatments: antimicrobial blue light (aBL), aBL combined with 5-aminolevulinic acid (aBL+ALA), rose bengal (RB), new methylene blue (NMB), and the cationic porphyrin TMPyP, applied under short- and prolonged-exposure conditions [12]. That study identified a common early stress response shared across all short treatments, and divergent, treatment-specific adaptive responses during prolonged exposure. However, the shared early response was characterized primarily at the level of gene-set overlap, leaving the relative magnitude, cross-treatment consistency, regulatory organization, and functional relevance of individual conserved-response genes unresolved.

Here, we focus specifically on this conserved early response. We hypothesized that chemically distinct photodynamic treatments would converge on a directionally stable transcriptional program containing genes that contribute to bacterial survival and post-treatment recovery. We prioritized conserved-response genes according to their response magnitude and cross-treatment consistency, characterized the regulatory architecture of this response, and assessed the functional contribution of selected genes using Keio deletion mutants and RT-qPCR validation. This integrated analysis links transcriptional conservation and regulatory organization to measurable survival and recovery phenotypes. Defining this conserved early survival program may identify stress-response pathways that could be targeted to enhance the efficacy of photodynamic methods.

## 2. Materials and Methods

### 2.1. Bacterial strains and culture conditions

RNA-seq and RT-qPCR experiments were performed using *Escherichia coli* BW25113 (parental strain originating from the *E. coli* Keio collection) under culture conditions previously described [12]. Briefly, transcriptomic experiments were conducted in M9 minimal medium supplemented as described in the original study.

Functional validation experiments were conducted using the parental BW25113 strain and selected single-gene knockout mutants from the Keio collection. For these assays, strains were cultured in Luria-Bertani (LB) broth or on LB agar plates containing 1.5% agar at 37°C under aerobic conditions. Knockout mutants were maintained in the presence of 15 µg/mL kanamycin. Strains were stored in 96-deep well plates filled with LB, and 15% glycerol was added to the medium for storage at −80°C. Before use, the cells were freshly stamped into new microtiter plates filled with LB medium and incubated overnight (16–20 h) at 37°C with shaking at 150 rpm.

### 2.2. RNA-seq data

The bacterial strain, sub-lethal photodynamic treatment parameters, RNA isolation, library preparation, and sequencing procedures were performed as previously described [12]. Briefly, *E. coli* BW25113 was subjected to five short-term (30 min) sub-lethal photodynamic treatments, including antimicrobial blue light (aBL), aBL combined with 5-aminolevulinic acid (aBL+ALA), rose bengal (RB) with green light, new methylene blue (NMB) with red light, and cationic porphyrin (TMPyP) with red light. Sub-lethal conditions were defined as those resulting in approximately a 1 log_10_ reduction in CFU/mL during the mid-log phase of growth over 30 minutes of exposure (detailed experimental conditions are presented in Supplementary Table S1). At least three independent biological replicates were analyzed per condition. RNA-seq libraries were prepared following rRNA depletion and sequenced on an Illumina NovaSeq X platform (paired-end, 150 bp). Raw data processing, quality control, mapping to the *E. coli* K-12 BW25113 reference genome (ASM584v2), read counting, and differential expression analysis were performed as previously described [12].

### 2.3. Bioinformatic and statistical analysis

To identify a modality-independent transcriptional response, a conserved short-term photodynamic core gene set was defined as the intersection of genes classified as differentially expressed in all five short-term treatments (aBL, aBL+ALA, RB, NMB, TMPyP), using identical statistical thresholds in each dataset (adjusted p-value < 0.05; fold change ≥ 1.5).

#### 2.3.1. Stratification of conserved core genes

To resolve heterogeneity within the intersection-defined conserved core, genes were stratified using two complementary measures: pooled response magnitude and cross-treatment consistency. For each gene, treatment-specific DESeq2 log₂ fold-change estimates and their standard errors were combined using DerSimonian-Laird random-effects meta-analysis with inverse-variance weighting [13]. This yielded a pooled log₂ fold change, a meta-analytic p-value, and heterogeneity statistics, including Cochran’s Q, τ², and I². P-values were adjusted across genes using the Benjamini-Hochberg procedure. Heterogeneity statistics were used descriptively to quantify variation in response amplitude across treatments. Cross-treatment consistency was assessed using robust rank aggregation [14]. Within each treatment, genes were ranked by log₂ fold change separately for induced and repressed responses, and RRA scores were then assigned to conserved-core genes according to their consensus direction. RRA was used to identify genes consistently ranked among the strongest responders across treatments. Directional concordance was defined as the fraction of treatments in which a gene changed in the consensus direction. Genes were classified as high-confidence core genes if they met all of the following criteria: meta-analysis FDR < 0.05, |pooled log₂FC| ≥ log₂(1.5), concordant direction of change in all five treatments, RRA FDR < 0.05, and membership in the top decile after averaging the meta-analysis and RRA-based ranks. Genes that were significant and directionally concordant but did not meet all high-confidence criteria were assigned to the intermediate tier, whereas genes failing the significance or direction-concordance criteria were classified as peripheral.

#### 2.3.2. Heatmap visualization of the conserved core

To visualize the dominant transcriptional structure within the conserved photodynamic core, the 25 highest-priority upregulated and 25 highest-priority downregulated genes were selected from the conserved core ranking. Gene priority was defined as the averaged rank derived from the random-effects meta-analysis and robust rank aggregation described above, with lower values indicating higher priority. For each selected gene, treatment-specific DESeq2 log₂ fold-change estimates relative to untreated controls were extracted and arranged into a gene-by-treatment matrix. Genes were grouped by consensus direction of regulation and ordered by priority within each group. Treatments were clustered hierarchically based on their log₂ fold-change profiles across the selected genes. Heatmaps were generated using the pheatmap package in R with a divergent color scale centered on zero. Functional and regulatory module annotations were assigned manually based on known gene functions and regulatory associations.

#### 2.3.3. Regulon enrichment and regulatory network analysis

Regulatory enrichment of the conserved photodynamic core was assessed using RegulonDB release 14.5. Transcription-factor and sigma-factor regulator-target interactions were extracted from the RegulonDB regulator-gene and sigma-gene network files and used to test upregulated and downregulated core genes separately. Enrichment was calculated for each regulon using a one-sided Fisher’s exact test, with RegulonDB-annotated genes as the background. Regulons with fewer than five annotated targets in the background were excluded, and p-values were adjusted using the Benjamini-Hochberg procedure. For each regulon, the number of conserved-core target genes, total regulon size, fold enrichment, nominal p-value, and FDR were reported in Table S3.

### 2.4. Photodynamic treatment and growth analysis of Keio mutants

Overnight cultures of the parental strain and selected knockout mutants prepared in microtiter plates were diluted fiftyfold in fresh LB medium in a new microtiter plate. For aBL and aBL+ALA studies, 200 µL of cultures were immediately subjected to irradiation, whereas for aPDI studies, 20 µL of photosensitizer was added to obtain the desired concentration in the final 200 µL volume, and the cultures were incubated in the dark for 15 min. After that, cultures were subjected to illumination. The detailed sub-lethal conditions for each treatment are provided in the Supplementary Table S1. For this study, new sub-lethal doses had to be estimated because knock-out mutants were grown in LB medium supplemented with kanamycin, whereas RNA-seq was performed in M9 medium. After illumination, plates were immediately placed in an Epoch 2 microplate reader (BioTek, USA), and the optical density (OD_600_) was measured every 30 min for 20 h at 37°C. The experiment was performed in three independent biological replicates.

### 2.5. Primer design and RT-qPCR

Culture preparation for RNA isolation for RT-qPCR analysis was performed as described for RNA-seq experiments. Briefly, overnight cultures of *E. coli* were diluted 1:25 into 10 mL of fresh, supplemented M9 medium and incubated for approximately 5.5 h at 37°C with shaking (150 rpm) to reach the exponential growth phase. Cultures were then adjusted to an OD600 of 0.3 and treated with sublethal doses of aBL/aPDI, as previously determined for RNA-seq analyses (Supplementary Table S1). Immediately after illumination, samples were incubated for 40 min at 37°C, after which cells were harvested directly into RNA Protect Bacteria Reagent (Qiagen, Germany) to stabilize RNA. Total RNA was purified using the RNeasy^®^ Mini Kit (Qiagen, Germany), following the manufacturer’s instructions. RNA concentration and purity were assessed using a NanoDrop One spectrophotometer (Thermo Scientific, USA), and samples were stored at −80 °C until further processing. Genomic DNA elimination and reverse transcription were performed using the QuantiTect Reverse Transcription Kit (Qiagen, Germany) according to the manufacturer’s instructions. Relative expression changes of *cpxP*, *degP*, *entB*, *fepA*, *gatA*, *hdeB*, *hslO*, *ibpB*, *modA*, *soxS*, and *spy* were analyzed using *csdA* and *bepA* as reference genes. The reference genes were selected based on their stable expression across all RNA-seq treatment and control datasets. RT-qPCR analysis was performed using three independent biological replicates, each analyzed in three technical replicates. Relative fold changes were calculated using the 2^−ΔΔCt^ method and log2-transformed for comparison with RNA-seq-derived expression changes. Primer sequences and qPCR conditions are provided in the Supplementary Table S4 and S5.

### 2.6. Statistical analysis

Differential expression was analyzed using DESeq2 v1.34.0, and genes with a fold change ≥ 1.5 and a Benjamini-Hochberg-adjusted p-value < 0.05 were considered differentially expressed. Treatment-specific log₂ fold changes were integrated using DerSimonian-Laird random-effects meta-analysis and robust rank aggregation, with Benjamini-Hochberg correction for multiple testing. Regulon enrichment was assessed using one-sided Fisher’s exact tests with Benjamini-Hochberg correction, restricting the analysis to regulons with at least five annotated targets. Phenotypic data are presented as mean ± standard deviation from three independent biological replicates. For each photodynamic treatment, knockout mutants were compared with the parental BW25113 strain using Dunnett’s multiple-comparison test. Agreement between RNA-seq and RT-qPCR log₂ fold-change values was assessed using Pearson’s correlation coefficient. A p-value or adjusted p-value < 0.05 was considered statistically significant. Analyses were performed in R.

## 3. Results

### 3.1. Identification of a conserved transcriptional response to short-term photodynamic stress

To identify genes consistently responding to photodynamic stress, differentially expressed genes (fold change ≥ 1.5; adjusted p-value < 0.05) from five short-term photodynamic treatments (aBL, aBL+ALA, RB, NMB, and TMPyP) were integrated using an intersection-based approach **(Fig. 1A)**. Genes that were differentially expressed across all five conditions were defined as the conserved transcriptional core. This analysis identified 891 genes shared across all treatments, indicating a substantial transcriptional program common to multiple photodynamic modalities. To further examine the structure of this conserved response, genes were classified by the direction of regulation across treatments **(Fig. 1B)**. Among the conserved genes, 500 (56.1%) were consistently upregulated, whereas 374 (42%) were consistently downregulated across all five conditions. Only 17 genes (<2%) showed variable regulation across treatments. This high level of directional consistency suggests that short-term photodynamic stress triggers a conserved regulatory response in *E. coli* despite differences in photosensitizer chemistry and light wavelength.

**Figure 1.**
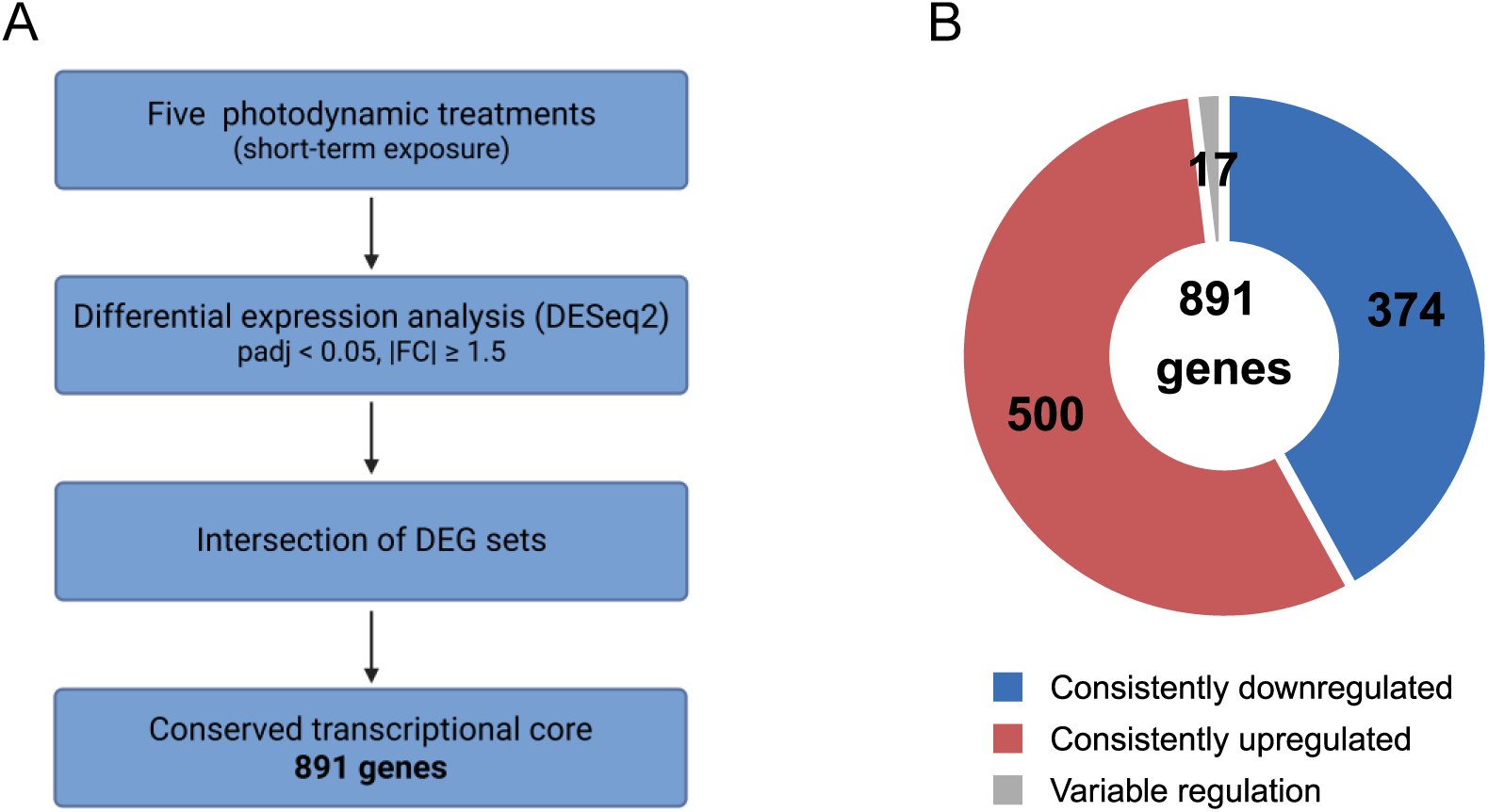
Identification of the conserved photodynamic transcriptional core in *E. coli* BW25113. (A) Workflow illustrating the identification of genes consistently differentially expressed across five short-term photodynamic treatments. **(B)** Architecture of the conserved transcriptional response showing the proportion of genes consistently upregulated (red), downregulated (blue), or displaying mixed regulation patterns (gray).

### 3.2. The conserved core is directionally stable but varies in amplitude

The five-treatment intersection identified 891 conserved core genes. To distinguish the most robust component of this response from genes that were consistently differentially expressed but less strongly prioritized, we stratified the core along two complementary axes: pooled effect size from a random-effects (DerSimonian-Laird) meta-analysis of the treatment-specific DESeq2 log₂ fold changes, and cross-treatment consistency from robust rank aggregation (RRA) of the treatment-level gene rankings **(Fig. 2A)**. Genes were ranked by absolute pooled effect size and RRA score, and the two ranks were averaged into a conserved response score; RRA significance is shown in Fig. 2A as −log₁₀ RRA FDR. The core was highly directionally conserved: 874 of 891 genes, corresponding to approximately 98%, changed in the same direction across all five treatments. Response amplitude, by contrast, varied substantially (median I² = 84%; **Fig. 2B),** indicating that heterogeneity primarily reflected differences in response strength rather than regulatory reversals. Finally, the integrated prioritization separated the core into 88 high-confidence genes that responded strongly and consistently across all treatments, and a larger intermediate set of 786 directionally concordant genes. The remaining 17 genes were classified as peripheral and corresponded to the genes showing variable directions of regulation across treatments, as described above (**Table S2**). The high-confidence set included 70 induced and 18 repressed genes. Induced genes were dominated by envelope stress, protein quality control, and oxidative or multidrug defense functions. Repressed genes included acid resistance, stationary-phase, and anaerobic metabolism programs. Together, these results define a directionally stable conserved response characterized by activation of envelope, proteotoxic, and oxidative-defense pathways and repression of acid-resistance and anaerobic-survival functions.

**Figure 2.**
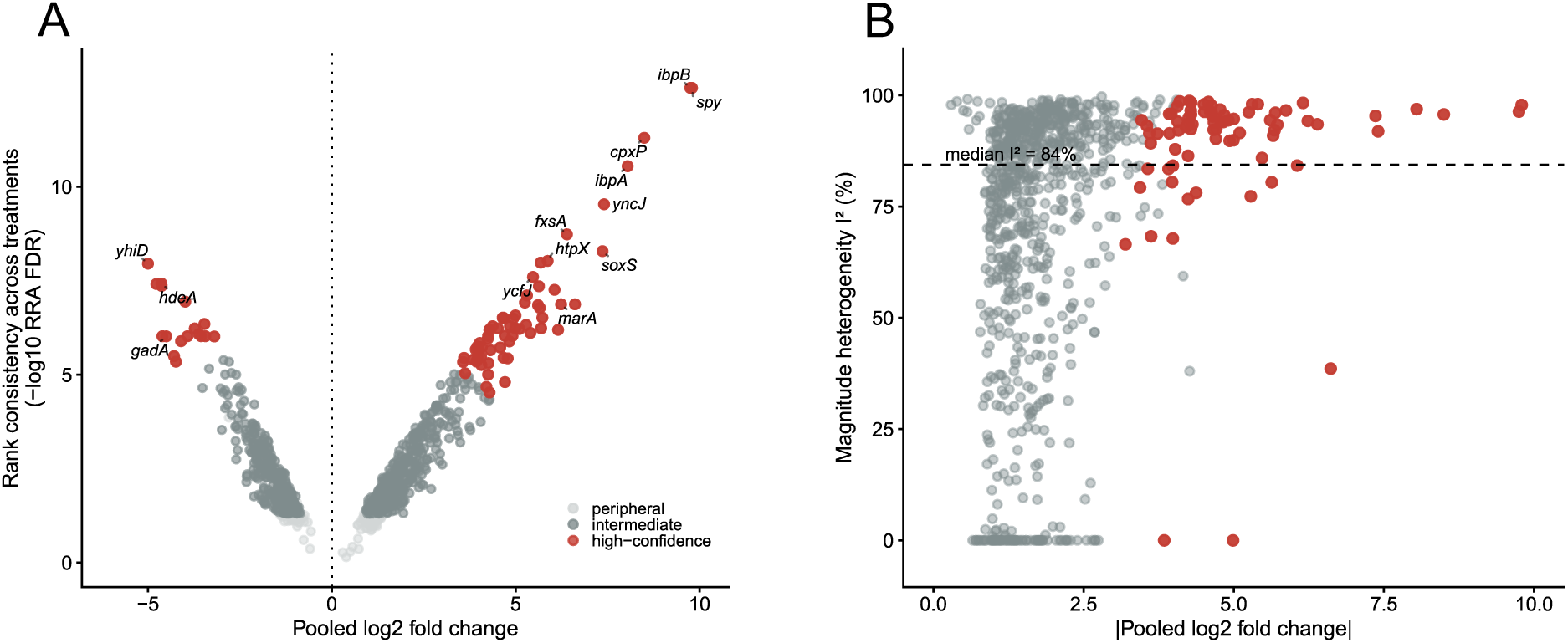
Stratification of the conserved photodynamic core in *E. coli* BW25113. The conserved core comprised 891 genes differentially expressed in all five short-term treatments; |log₂FC| ≥ log₂(1.5), adjusted *p* < 0.05 in each comparison. **(A)** Core genes were prioritized by pooled effect size from DerSimonian-Laird random-effects meta-analysis and by cross-treatment rank consistency assessed with robust rank aggregation (RRA). The y-axis shows −log₁₀ RRA FDR; higher values indicate genes consistently ranked among the strongest responders across treatments. Selected high-confidence genes are labeled. **(B)** Magnitude heterogeneity, expressed as I², plotted against absolute pooled log₂ fold change. The dashed line indicates the median I² across core genes. High-confidence genes are shown in red. High-confidence genes met all prioritization criteria: meta-analysis FDR < 0.05, |pooled log₂FC| ≥ log₂(1.5), RRA FDR < 0.05, concordant direction in all five treatments, and top-decile combined priority.

### 3.3. A conserved transcriptional program underlies the response to photodynamic stress

To identify the dominant component of the conserved photodynamic response, we selected the 25 highest-ranked upregulated and 25 highest-ranked downregulated genes by conserved response score and visualized their treatment-specific log₂ fold changes **(Fig. 3)**. These genes showed a highly consistent directionality across all five conditions, although response magnitude varied between photosensitizers. Rose bengal produced overall weaker changes within this prioritized set than the porphyrin- and phenothiazinium-based treatments, consistent with the amplitude differences observed in the conserved-core ranking. The upregulated genes were dominated by envelope stress and protein quality control functions, i.e., the periplasmic chaperone *spy*, Cpx-associated genes such as *cpxP* and *yebE*, the small heat-shock proteins *ibpA* and *ibpB*, the membrane protease *htpX*, and the major chaperone *dnaK*. Additional induced genes pointed to oxidative and multidrug stress responses, including members of the sox and mar regulons, as well as genes linked to phage-shock, phosphate-starvation, and efflux functions. In contrast, the downregulated genes were enriched for acid resistance, anaerobic metabolism, and stationary-phase-associated functions, including the glutamate-dependent acid-resistance system, acid-stress chaperones, hydrogenase-associated genes, the molybdate transport operon, and curli-biogenesis genes. Thus, despite differences in photosensitizer chemistry and activating wavelength, the five treatments converged on a shared transcriptional program: induction of envelope, oxidative, and proteotoxic defenses together with repression of acid-resistance, anaerobic, and stationary-phase survival functions.

**Figure 3.**
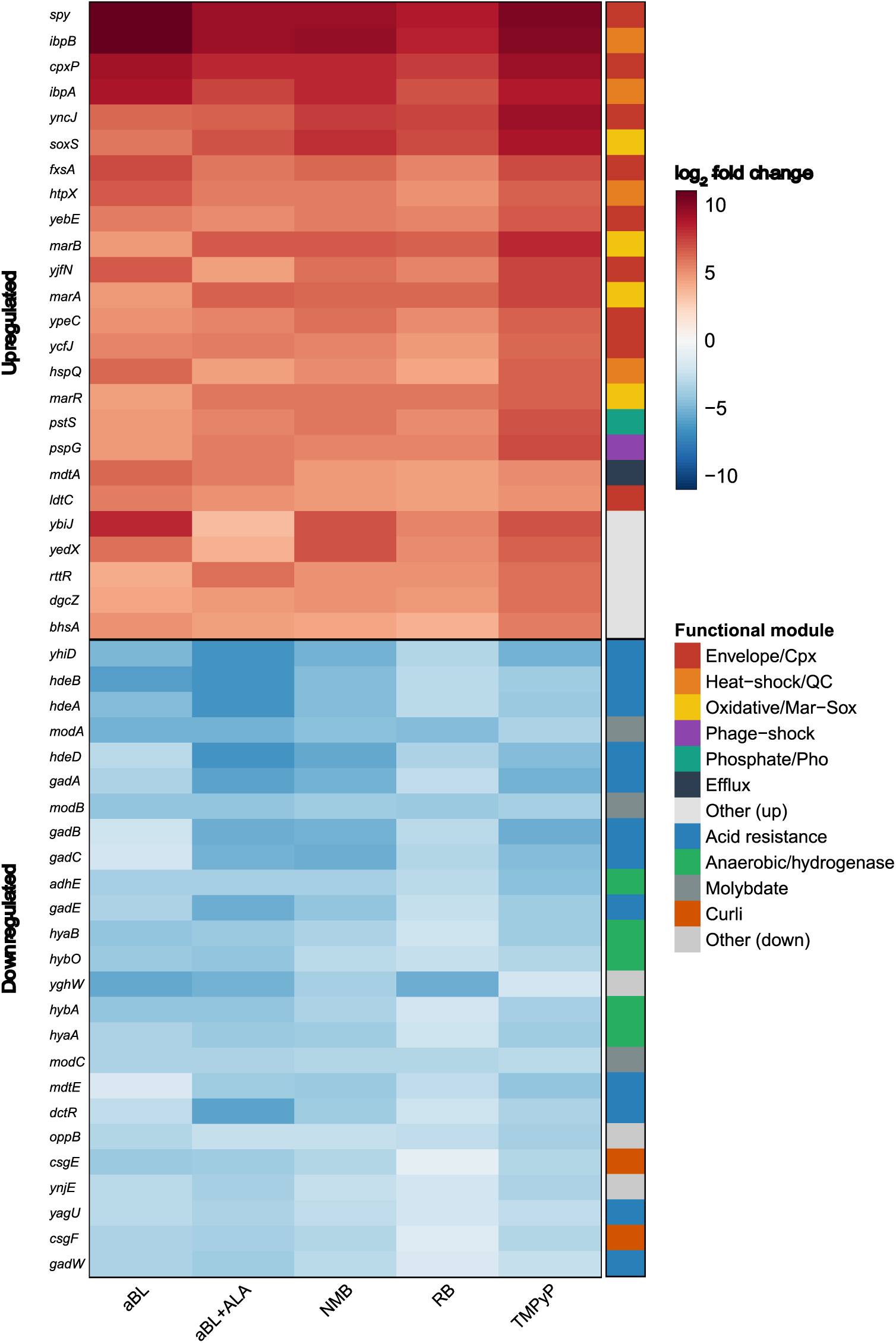
A conserved core transcriptional program defines the response to photodynamic stress in *E. coli* BW25113. Heatmap of treatment-specific log₂ fold changes for the 25 highest-priority upregulated and 25 highest-priority downregulated genes, ranked by the conserved response. Rows represent genes and columns represent photodynamic treatments. Functional annotations highlight major stress-response categories represented among the prioritized genes. Data are from at least three independent biological replicates per treatment.

### 3.4. The conserved response is organized around heat-shock, envelope-stress and acid-resistance regulators

Regulon enrichment analysis of the conserved core revealed a compact regulatory architecture associated with the photodynamic stress response. Upregulated and downregulated core gene sets were tested separately for enrichment of RegulonDB transcription-factor and sigma-factor regulons, using RegulonDB-annotated genes as the background and retaining regulons with at least five annotated targets. Among the induced core, the most extensive and statistically significant enrichment was observed for the heat-shock sigma factor σ^32^ (RpoH) (58 core genes, ∼2.8-fold enrichment, FDR ≈ 4 × 10⁻¹²), accounting for the protein-quality-control module identified in Fig. 3, including *ibpA*, *ibpB*, *dnaK*, *groES,* and *htpX*. In parallel, the induced core was enriched for envelope-stress regulons, including CpxR, BaeR, the extracytoplasmic sigma factor σ^24^ (RpoE), the phage-shock regulator PspF, and the LysR-type regulator YhaJ (**Fig. 4A, C**). Thus, the induced conserved response converges on two complementary regulatory axes: cytoplasmic protein quality control, centered on σ^32^, and cell-envelope stress, associated with CpxR, BaeR, σ^24^, and PspF. Although several oxidative- and multidrug-defense genes (e.g., *soxS*, *marA*) are present in the conserved core (Fig. 3), no corresponding regulon (OxyR, SoxS, MarA, Rob) was significantly enriched, indicating individual-gene rather than regulon-level induction. The downregulated core was enriched for regulators of acid resistance, metal and iron homeostasis, and stationary-phase physiology (**Fig. 4B**). The acid-resistance signal was dominated by the glutamate-dependent system, with enrichment of GadW, GadX, and GadE regulons, consistent with repression of *gad* genes in the conserved core. Other enriched regulons included ModE and Fur, together with σ^38^ (RpoS), ArcA, and H-NS. The complete enrichment results are provided in **Table S3**. Overall, the conserved photodynamic response links induction of protein-quality-control and envelope-stress pathways with repression of acid-resistance, metal- and iron-associated, anaerobic, and stationary-phase-associated functions.

**Figure 4.**
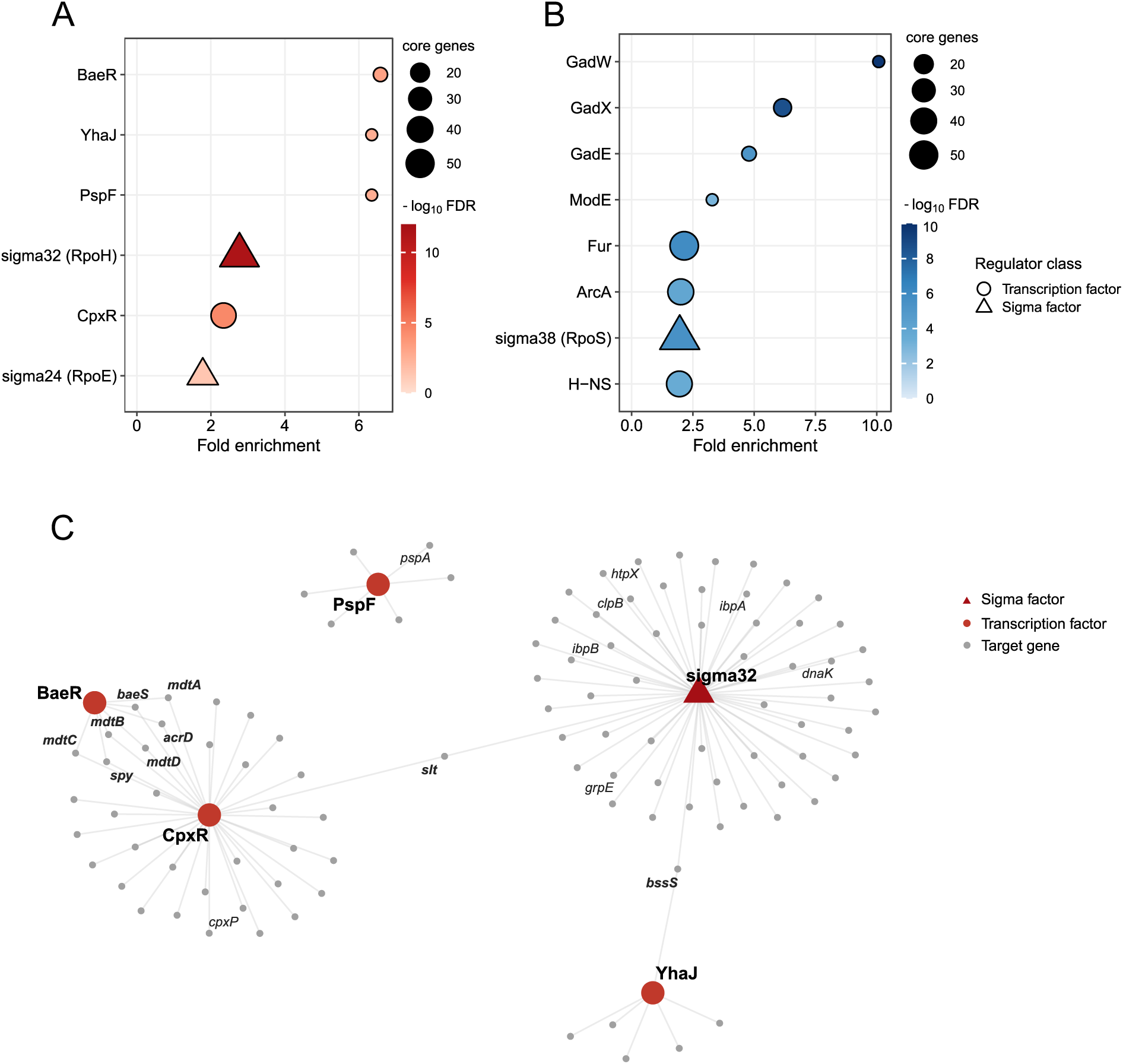
Regulon enrichment and regulatory network architecture of the conserved photodynamic response. Regulon enrichment analysis was performed for genes consistently upregulated **(A)** or downregulated **(B)** across treatments. Enrichment was calculated relative to RegulonDB-annotated genes using one-sided Fisher’s exact tests with Benjamini–Hochberg correction. Point size indicates the number of conserved-core genes assigned to each regulon, and color indicates statistical significance (−log₁₀ FDR). On graphs, only representative enriched regulators are shown. **(C)** Regulatory network of selected enriched regulators and their target genes within the upregulated conserved core. Regulator nodes are shown separately from target genes; edges represent curated RegulonDB regulator-target interactions. Genes connected to at least two regulators are bolded.

### 3.5. Conserved-core genes are functionally required for survival of photodynamic stress

To test whether the conserved transcriptional core reflects a functionally protective response, we selected seven consistently induced core genes representing the main upregulated modules: envelope stress (*spy*, *cpxP*), cytoplasmic protein quality control (*ibpB*, *htpX*), oxidative or multidrug defense (*soxS*, *grxA*), and the Psp inner-membrane stress response (*pspA*). The corresponding Keio single-gene deletion mutants and the parental strain were exposed to the five photodynamic treatments, and their responses were assessed by clonogenic survival, expressed as log₁₀ CFU reduction, and post-treatment growth, measured by OD₆₀₀. For growth, we quantified the integrated growth response as the area under the OD₆₀₀ growth curve (AUC), normalized to each strain’s own untreated control, and the recovery delay defined as the time required to reach ΔOD₆₀₀ = 0.2. In Fig. 5A and 5B, values are expressed relative to the parental strain; red therefore indicates increased mutant susceptibility relative to WT, whereas blue indicates reduced susceptibility or improved mutant performance relative to WT.

**Figure 5.**
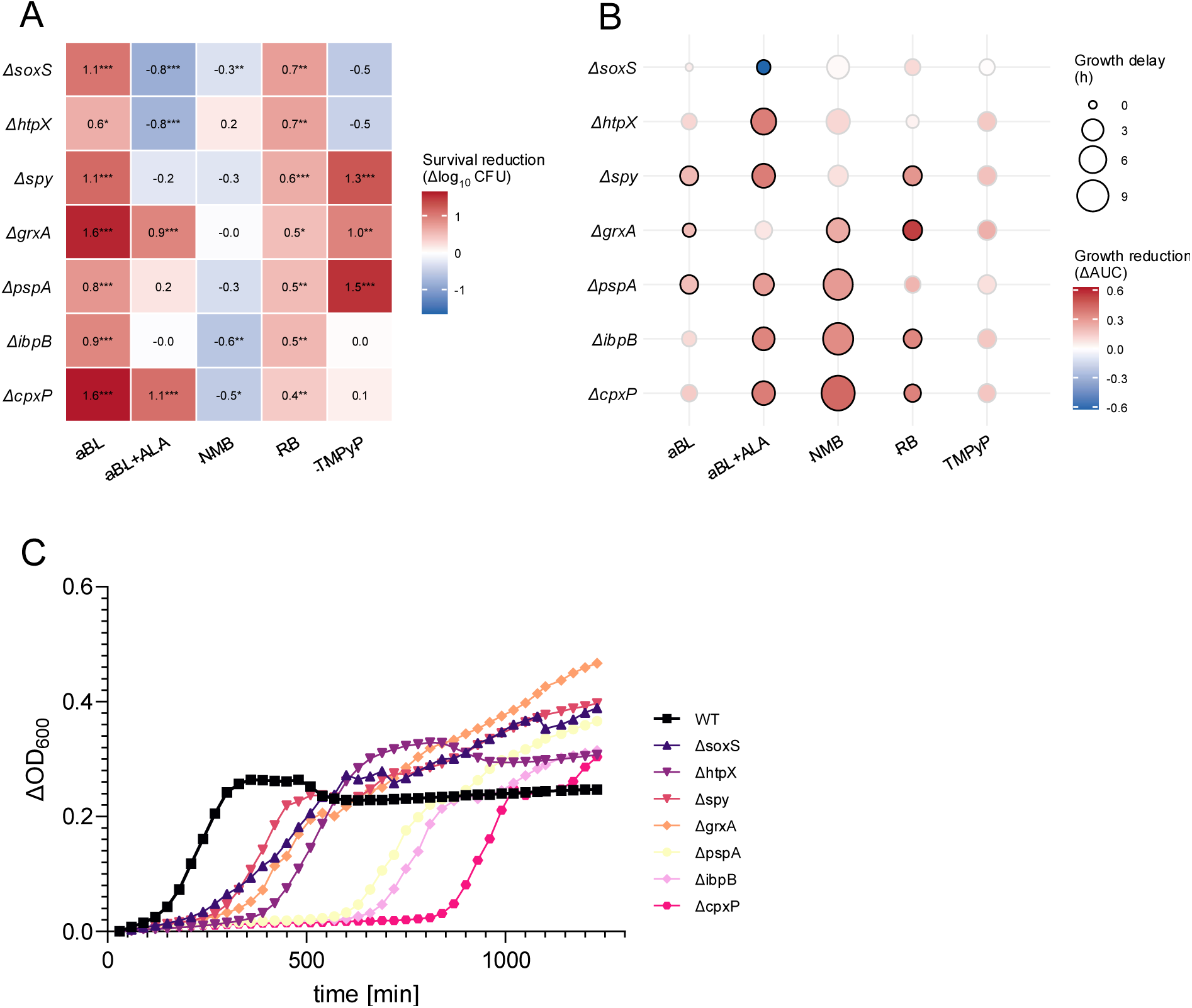
Conserved-core genes contribute to survival and recovery after photodynamic stress. Seven consistently induced high-confidence core genes were selected for functional validation using single-gene Keio deletion mutants: Δ*spy*, Δ*cpxP*, Δ*ibpB*, Δ*htpX*, Δ*soxS*, Δ*grxA* and Δ*pspA*. Mutants and the parental strain were exposed to aBL, aBL+ALA, NMB, RB, and TMPyP. **(A)** Survival sensitization, shown as the difference in log₁₀ CFU reduction between each mutant and the parental strain. **(B)** Growth-based sensitization, shown as growth reduction relative to WT. Dot color indicates the difference in growth retention between WT and mutant, calculated from the area under the OD₆₀₀ growth curve normalized to each strain’s untreated control. Dot size indicates recovery delay relative to WT, defined as the delay in reaching ΔOD₆₀₀ = 0.2. In panels A and B, red indicates increased mutant susceptibility relative to WT, whereas blue indicates reduced susceptibility or improved mutant performance relative to WT. Black outlines in panel B indicate significant differences from WT by Dunnett’s test (p < 0.05). **(C)** Representative post-treatment growth curves for tested strains under NMB treatment.

Deletion of conserved-core genes increased susceptibility to photodynamic killing in a treatment-dependent manner (**Fig. 5A**). The strongest and most consistent sensitization was observed under antimicrobial blue light, where all seven mutants showed greater log₁₀ CFU reduction than the parental strain. The largest effects were detected for Δ*cpxP* and Δ*grxA*, followed by Δ*soxS*, Δ*spy*, Δ*ibpB* and Δ*pspA*. Rose bengal treatment also sensitized all mutants, although the magnitude of the effect was generally lower than under aBL. TMPyP produced a more selective phenotype, with pronounced sensitization of Δ*spy*, Δ*grxA* and Δ*pspA*. In contrast, aBL combined with ALA and NMB showed more heterogeneous effects, including cases in which mutant survival was comparable to or higher than that of the parental strain. Thus, although the conserved response is shared transcriptionally across treatments, the contribution of individual genes to acute killing depends on the photodynamic modality. Growth-based phenotyping revealed an additional recovery component of mutant susceptibility (**Fig. 5B**). Several mutants displayed reduced growth retention relative to the parental strain and/or delayed post-treatment recovery, particularly after NMB, RB and aBL+ALA exposure. The dot-heatmap indicated that growth sensitization was often expressed as delayed resumption of growth rather than complete failure to reach high optical density. This was especially evident for mutants affecting envelope stress, protein quality control and oxidative defense, supporting a role for these modules in recovery after sub-lethal photodynamic injury. Significant growth phenotypes were detected for selected mutant– treatment combinations, as indicated by black outlines. Representative post-treatment growth curves are shown in Fig. 5C for NMB-treated cultures, illustrating the delayed recovery phenotype observed in selected mutants. Complete growth curves for all strains and photodynamic treatments are provided in the Supplementary Materials (**Fig. S1**).

The survival and growth assays captured different aspects of the mutant phenotype. The survival assay measured acute killing, whereas the growth assay measured the ability of surviving cells to recover after treatment. This distinction helped explain modality-specific patterns. Under aBL, most mutants showed increased killing, but their post-treatment growth was only modestly affected, suggesting that aBL-induced vulnerability was best captured by viability. In contrast, under NMB, several mutants showed delayed recovery without a corresponding increase in killing, indicating a mainly sub-lethal recovery defect. The aBL+ALA response was more heterogeneous: Δ*cpxP* and Δ*grxA* were sensitized, whereas Δ*soxS* showed higher growth retention than the parental strain and reduced susceptibility in the survival assay. Overall, these patterns suggest that the conserved response is modular, with different protective genes becoming important under different photodynamic modalities.

### 3.6. Validation of RNA-Seq data using RT-qPCR

To validate the RNA-seq results, 11 representative genes from the conserved response were selected for RT-qPCR analysis. These genes covered multiple functional categories, including envelope stress, protein quality control, oxidative stress regulation, iron acquisition, molybdate transport, acid stress, and carbohydrate metabolism. RT-qPCR confirmed the direction of expression changes for all analyzed gene-treatment combinations, with complete concordance across 55 comparisons. Moreover, RNA-seq and RT-qPCR log_2_ fold-change values were strongly correlated across the full validation dataset (Pearson r = 0.988, Fig. 6), supporting the robustness of the transcriptomic dataset. The full validation dataset is provided in Table 1.

**Figure 6.**
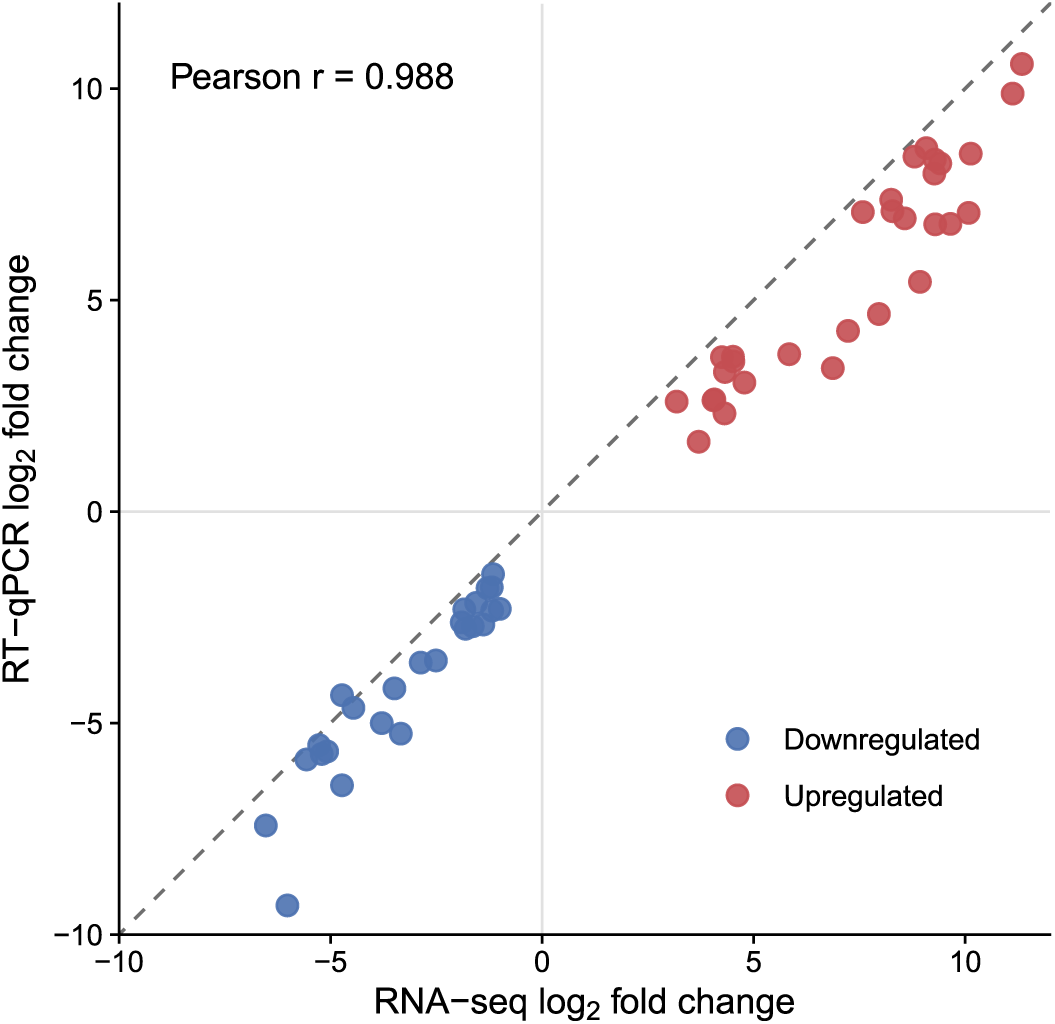
Validation of RNA-seq results by RT-qPCR. Scatter plot showing the relationship between RNA-seq-derived and RT-qPCR-derived log₂ fold-change values for 11 selected conserved-response genes across five photodynamic treatments. Each point represents one gene-treatment comparison. Red and blue points indicate induced and repressed genes, respectively. The dashed line indicates equal log₂ fold-change values between both methods. Pearson correlation was calculated across all 55 comparisons.

**Table 1.**
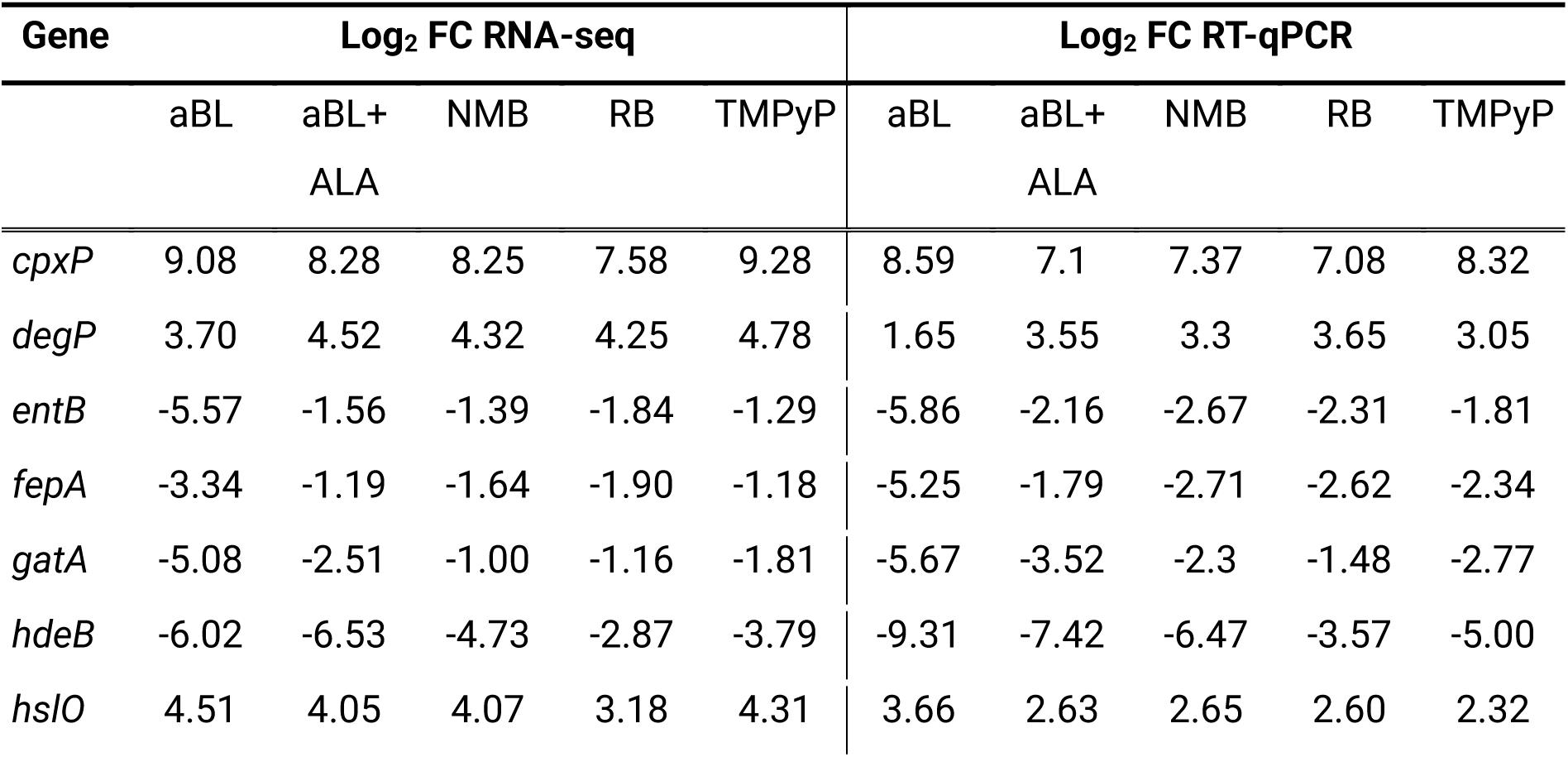
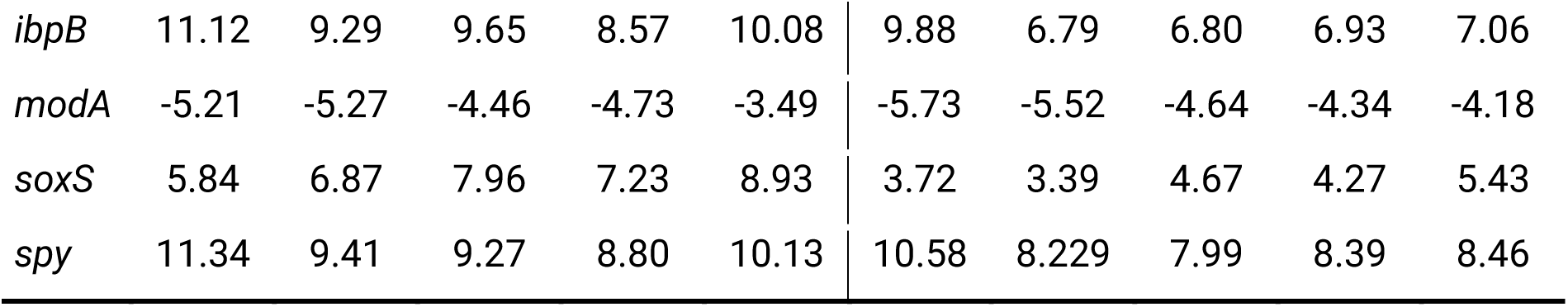
Comparison of RNA-seq- and RT-qPCR-derived log₂ fold-change values for 11 selected conserved-response genes across five photodynamic treatments.

| Gene | Log<sub>2</sub> FC RNA-seq |  |  |  |  | Log<sub>2</sub> FC RT-qPCR |  |  |  |  |
| --- | --- | --- | --- | --- | --- | --- | --- | --- | --- | --- |
|  | aBL | aBL+<br>ALA | NMB | RB | TMPyP | aBL | aBL+<br>ALA | NMB | RB | TMPyP |
| *cpxP* | 9.08 | 8.28 | 8.25 | 7.58 | 9.28 | 8.59 | 7.1 | 7.37 | 7.08 | 8.32 |
| *degP* | 3.70 | 4.52 | 4.32 | 4.25 | 4.78 | 1.65 | 3.55 | 3.3 | 3.65 | 3.05 |
| *entB* | -5.57 | -1.56 | -1.39 | -1.84 | -1.29 | -5.86 | -2.16 | -2.67 | -2.31 | -1.81 |
| *fepA* | -3.34 | -1.19 | -1.64 | -1.90 | -1.18 | -5.25 | -1.79 | -2.71 | -2.62 | -2.34 |
| *gatA* | -5.08 | -2.51 | -1.00 | -1.16 | -1.81 | -5.67 | -3.52 | -2.3 | -1.48 | -2.77 |
| *hdeB* | -6.02 | -6.53 | -4.73 | -2.87 | -3.79 | -9.31 | -7.42 | -6.47 | -3.57 | -5.00 |
| *hsIO* | 4.51 | 4.05 | 4.07 | 3.18 | 4.31 | 3.66 | 2.63 | 2.65 | 2.60 | 2.32 |
| *ibpB* | 11.12 | 9.29 | 9.65 | 8.57 | 10.08 | 9.88 | 6.79 | 6.80 | 6.93 | 7.06 |
| *modA* | -5.21 | -5.27 | -4.46 | -4.73 | -3.49 | -5.73 | -5.52 | -4.64 | -4.34 | -4.18 |
| *soxS* | 5.84 | 6.87 | 7.96 | 7.23 | 8.93 | 3.72 | 3.39 | 4.67 | 4.27 | 5.43 |
| *spy* | 11.34 | 9.41 | 9.27 | 8.80 | 10.13 | 10.58 | 8.229 | 7.99 | 8.39 | 8.46 |

## 4. Discussion

The current study utilized time-resolved global transcriptomics to define the systems-level physiological response of *Escherichia coli* BW25113 to five distinct sublethal phototreatments: antimicrobial blue light (aBL), aBL with 5-aminolevulinic acid (aBL+ALA), rose bengal (RB), new methylene blue (NMB), and the cationic porphyrin (TMPyP). Intersection analysis identified a unified, conserved early survival program consisting of 891 shared differentially expressed genes (DEGs). Remarkably, 98% of these genes exhibited consistent directionality across all treatments, despite fundamental differences in photosensitizer chemistry and activating wavelengths (Fig. 1).

Using meta-analysis and robust rank aggregation, we prioritized 88 high-confidence core genes (Fig. 2). The dominant component of this conserved response was characterized by a massive induction of the envelope stress response (ESR) and cytoplasmic protein quality control pathways, alongside a systemic repression of acid resistance (the *gad* system), hydrogen metabolism, and biofilm formation modules (Fig. 3). Regulon enrichment identified that induced genes were primarily organized around the heat-shock sigma factor σ³²/RpoH and the envelope-stress regulators CpxR, BaeR, σ²⁴/RpoE, and PspF (Fig. 4). Functional validation using Keio collection mutants (e.g., Δ*spy*, Δ*cpxP*, Δ*ibpB*, and Δ*htpX*) confirmed that these core genes are critical for survival and post-treatment recovery, as their deletion significantly sensitized *E. coli* to photodynamic challenge (Fig. 5).

Our investigation demonstrates that chemically diverse photodynamic treatments converge on a conserved early survival program in *E. coli*. This finding shifts the focus from photosensitizer-specific mechanisms to a universal bacterial response centered on envelope integrity and protein homeostasis during the initial stages of stress.

The discovery of this unified 891-gene core defines the acute stress architecture that bacteria recruit within the first minutes of photodynamic challenge. Our previously published data [12] demonstrated that this massive transcriptional remodeling, which affects up to 58% of the genome, is characteristic of short-term exposure (30 min). Interestingly, we have evidenced that as the duration of stress increases to 7–8 hours, the bacterial response transitions from this unified core toward highly treatment-specific adaptive trajectories. This transition supports a two-phase adaptation model, i.e., an early phase of redox defense followed by a late phase of resource conservation and metabolic reprogramming.

The conserved core response is anchored by the massive induction of envelope stress regulons, particularly CpxR, BaeR, and σ²⁴ (RpoE) (Fig. 4A). A study by Chittò et al. supported our results and demonstrated that while different photosensitizers have distinct primary sub-compartment targets such as methylene blue (MB) targeting the cytosol and silicon phthalocyanine (SiPc) targeting the envelope, they ultimately trigger specialized stress-response modules to maintain envelope homeostasis [15]. Similarly, Ziegelhoffer and Donohue evidenced that light-dependent processes generate singlet oxygen (¹O₂), which necessitates the recruitment of extracytoplasmic function sigma factors like σ^E^ (RpoE) to repair membrane damage [16]. Our data further confirm this by showing that deletion of core envelope stress response genes like *spy* and *cpxP* results in significant survival sensitization and recovery delays across all tested phototreatments (Fig. 5A, B).

Beyond membrane stress, our findings highlight a profound enrichment of the σ³² (RpoH) regulon, managing cytoplasmic protein quality control (Fig. 4A). The induction of high-priority core genes such as *dnaK*, *ibpA/B*, and *groES* (Fig. 3) indicates that sublethal aPDI causes substantial proteotoxic stress. A study by Imlay may be supportive of these results, as it described how ROS collide randomly with enzymes and remove electrons, which destroys their activity, leading to the destruction of metalloenzymes and the formation of protein carbonyls [17]. Moreover, Manoil et al. reported a similar up-regulation of *dnaK* in *Enterococcus faecalis* following aPDI, suggesting that protein refolding and recycling are universally critical for surviving a photodynamic burst [18]. Our functional data support this, as the Δ*ibpB* and Δ*htpX* mutants exhibited diminished growth retention and significant recovery delays (Fig. 5B).

In addition, the conserved core response involves a systemic metabolic downshift, including the repression of oxidative phosphorylation and amino acid biosynthesis, i.e., the *mod* operon or the *gad* system (Fig. 3). In our previously published data [12], we have described this as a protective mechanism to limit endogenous ROS production from the electron transport chain during the acute phase of stress. Rapacka-Zdonczyk demonstrated that such states are often associated with reduced growth and metabolic activity, which can lead to transient non-genetic tolerance or persistence. The observed metabolic downshift is further supported by the systemic repression of the RpoS regulon during the acute (30 min) phase of the photodynamic stress response (Fig. 4B). While RpoS is traditionally defined as the master regulator of the stationary phase and general stress response, Adams et al. evidenced that the temporal framework for the transcription of RpoS-dependent genes varies significantly depending on the specific environmental challenge [19]. In this context, the early repression of the RpoS regulon observed in our study suggests that when faced with a rapid photodynamic burst, the bacterial cell prioritizes specialized local repair mechanisms such as the RpoH (heat-shock) and RpoE (envelope stress) pathways while temporarily suspending the global physiological reorganization typically governed by RpoS to conserve energy resources critical for immediate survival. This strategic delay in RpoS-mediated adaptation aligns with a two-phase model where the cell first focuses on immediate redox defense and protein/envelope stabilization before transitioning to a more permanent adaptive state.

Next, an unexpected signature of the conserved core was the strong repression of the glutamate-dependent acid resistance (GDAR) system and biofilm-associated genes such as curli-specific genes (*csg*) (Fig. 3). Waterman and Small evidenced that GDAR genes are typically induced under stationary-phase or acidic conditions [20]. The systematic downregulation observed here suggests that acute photo-oxidative stress actively discourages energy-intensive survival strategies in favor of immediate cellular repair. In summary, the repression of GDAR and *csg* genes confirms that *E. coli* treats photodynamic stress as an emergency state, abandoning luxury survival programs in favor of the immediate maintenance of envelope integrity and protein homeostasis.

## 5. Conclusion

In conclusion, our study defines the acute stress architecture of *E. coli* under diverse photodynamic challenges. We have demonstrated that chemically distinct phototreatments converge on a conserved early survival program that prioritizes the maintenance of envelope integrity and cytoplasmic protein quality control. By integrating these findings with our previously published time-resolved data, we establish that this acute 891-gene response is the critical first phase of a two-phase adaptation model, which eventually diverges into treatment-specific metabolic restructuring. The functional necessity of these core modules, validated through Keio mutant sensitization, provides a mechanistic basis for the rational design of light-based therapeutic protocols that target the bacterial transcriptional core to bypass antimicrobial resistance.

## Acknowledgments

The authors thank the National BioResource Project (NBRP, NIG, Japan) for contributing to our work by providing us with the *E. coli* BW25113 parental strain and mutants.

## Availability of data

The data generated or analyzed during this study are included in this published article and in the supplemental material. RNA-seq data are deposited in the NCBI Gene Expression Omnibus (GEO) database under accession number GSE317397.

